# HYPERURICAEMIA DOES NOT INTERFERE WITH AORTOPATHY IN A MURINE MODEL OF MARFAN SYNDROME

**DOI:** 10.1101/2023.05.21.541606

**Authors:** Isaac Rodríguez-Rovira, Angela López-Sainz, Maria Encarnación Palomo-Buitrago, Belén Pérez, Francesc Jiménez-Altayó, Victoria Campuzano, Gustavo Egea

## Abstract

Redox stress is involved in the aortic aneurysm pathogenesis in Marfan syndrome (MFS). We recently reported that allopurinol, a recognised inhibitor of XOR, inhibited aortopathy in a mouse model of MFS acting as an antioxidant. Hyperuricaemia is ambiguously associated with cardiovascular injuries as uric acid (UA), having antioxidant or pro-oxidant properties depending on the concentration and accumulation site. We aimed to evaluate whether hyperuricaemia causes harm or relief in MFS aortopathy and cardiopathy pathogenesis. Two-month-old male wild-type (WT) and MFS mice (*Fbn1^C1041G/+^*) were injected intraperitoneally for several weeks with potassium oxonate (PO), an inhibitor of uricase, an enzyme that catabolises UA to allantoin. Plasma UA and allantoin levels were measured via several techniques, aortic root diameter and cardiac parameters by ultrasound imaging, aortic wall structure by histopathology, and pNRF2 levels by immunofluorescence. PO induced a significant increase in UA in blood plasma both in WT and MFS mice, reaching a peak at three and four months of age but decaying at six months. Hyperuricaemic MFS mice showed no change in the characteristic aortic aneurysm progression or aortic wall disarray evidenced by large elastic laminae ruptures. There were no changes in cardiac parameters or the redox stress-induced nuclear translocation of pNRF2 in the aortic tunica media. Altogether, the results suggest that hyperuricaemia interferes neither with aortopathy nor cardiopathy in MFS mice.

## 1. INTRODUCTION

Redox stress (RS) results from the chronic dysfunctional balance between the production and elimination of both reactive oxygen and nitrogen species (ROS and RNS; RONS). RS is largely involved in numerous pathological processes leading to the loss- or gain-of-function of lipids and proteins and changes in gene expression depending on the main subcellular location where it occurs [1,2]. With the accompanying cell signalling (growth factors, kinases), haemodynamics, and extracellular matrix alterations, RS is becoming increasingly clearly an aggravating driver of both thoracic and abdominal aortic aneurysms, including MFS syndrome (MFS) [3-9].

MFS is a genetic connective tissue disorder caused by mutations in the fibrillin-1 gene (*FBN1*) affecting cardiovascular, ocular, and musculoskeletal systems [10-14]. The aortic root aneurysm, with its usual subsequent dissection and rupture, is the principal cause of death in MFS patients and prophylactic surgical replacement is the main treatment option to increase life expectancy [15-17]. Nonetheless, current pharmacological treatments with beta-blockers (atenolol) and angiotensin II receptor type I (ART1) antagonists (losartan; LOS) have recently provided a minor, yet significant, improvement in aortopathy progression [18]. In any case, new complementary treatment strategies are necessary to achieve a substantial improvement in slowing or stopping aortopathy progression considering the variety of molecular mechanisms involved. It is at this point where anti-redox stress agents, such as resveratrol [19-21], allopurinol [22,23], and cobinamide [24] have recently appeared on the scene as potential new therapies.

We reported that upregulated NADPH (nicotinamide adenine dinucleotide phosphate) oxidase 4 (NOX4) aggravates aortic aneurysm progression in both in human and mouse MFS aortic samples, and other investigators have subsequently established specific signalling pathways mediating this injury [25-27]. Besides NOX4, nitric oxide synthase [28], mitochondria [29], endoplasmic reticulum [30-31], and xanthine oxidoreductase (XOR) are other important sources of ROS in the cardiovascular system [32]. XOR is a complex enzyme that, concomitantly with uric acid (UA) formation from purine metabolism, also forms ROS (superoxide and hydrogen peroxide) [33]. XOR also participates in MFS aortopathy [22, 34].

In humans and great apes, UA can abnormally accumulate in plasma and some tissues. However, in rodents, UA is quickly catabolised by uricase to allantoin, which is much more water-soluble than UA [35]. In humans, allantoin has been suggested as an oxidative stress biomarker because it is not produced metabolically [36]. Several epidemiological studies suggest a relationship between elevated serum UA and cardiovascular risk factors. However, this association remains controversial [37-41]. Thus, UA has a dual role in redox biopathology [36, 42,43]. On the one hand, it accounts for as much as 50% of the total antioxidant capacity of biological fluids in humans [44,45], and it was postulated that higher UA levels protect against some peroxynitrite and inflammatory-induced CNS diseases [46]. On the other, when UA accumulates in the cytoplasm or the acidic/hydrophobic milieu, it becomes a pro-oxidant, promoting RS [47-49].

UA was found in the aortic walls of human aneurysms and atherosclerotic plaque arteries [50], and a positive correlation between serum UA levels and aortic dilatation and dissection has been reported [23,51-54]. Nonetheless, epidemiologic and biochemical studies on UA formation have demonstrated that it is not only UA itself that leads to a worse prognosis and increased cardiovascular events but also ROS formed during XOR activity. Therefore, the resulting combined action of excessive UA and ROS formation along with enhanced XOR activity could significantly contribute to oxidative stress-linked endothelial dysfunction and heart failure, perhaps in aortopathies as well.

Interestingly, the LIFE (Losartan Intervention for Endpoint reduction) clinical trial demonstrated that LOS was superior to atenolol in reducing cardiovascular events and mortality in patients with hypertension [56]. This result was partially attributable to the intrinsic uricosuric effect of LOS via inhibition of the UA transporter URAT1 [57]. In addition, LOS normalised the aortic dilatation in MFS mice better than beta-blockers based on the rationale that LOS inhibited ATR1-mediated TGFβ hypersignalling [58]. Considering that Ang II activates endothelial XOR [59], it is possible that the reparatory effect of LOS on the aortic aneurysm in MFS mice might also be partially attributable to its uricosuric effect. Moreover, in the heart and circulatory system, UA also stimulates the production of ROS via the activation of TGFβ1 and NOX4 [60,61].

Therefore, considering all the above, it might be possible that UA participates, to some extent, in MFS aortopathy, aggravating it by acting as a pro-oxidant and/or promoting ROS production or, just the opposite, mitigating aneurysm progression by acting as an antioxidant. Nevertheless, knowing the broad pathological implication of increased plasma UA levels, we are inclined to think that UA’s effect might be more aggravating than mitigating. Then, this study aimed to evaluate the impact of experimentally induced hyperuricaemia in aortic aneurysm progression in a murine model of MFS. Whereas in the observed allopurinol-induced blockade of aortopathy, plasma UA levels remained unaltered, and its involvement was initially discarded, this cannot be definitive as, physiologically, UA in rodents is catabolised to allantoin by uricase. Therefore, to study our aim in MFS mice, it was necessary to generate and maintain increased plasma UA levels long enough to coincide with aneurysm formation to evaluate its progression. This transient hyperuricaemic model is feasible utilising uricase inhibitors like potassium oxonate (PO). Our results indicate that, despite the presence of high plasma UA levels, aortic aneurysm progression in MFS mice was not affected in any way (aggravating or ameliorating), indicating that UA does not mediate in murine Marfan aortopathy.

## 2. RESULTS

### 2.1. Sex differences in uric acid and allantoin blood plasma levels in MFS mice

We first measured plasma levels of UA and allantoin in two-month-old male and female WT and MFS mice, age of the basal start point of subsequent PO treatment (see supplementary Figure 1 for a scheme of experimental protocols followed in the study), and at which the aneurysm can already be clearly seen in both sexes (supplementary Figure 2), There were no differences in basal plasma UA levels between WT male and female. Unlike in male, there were significant differences between WT and MFS female as well as between MFS male and female being significantly higher in the former (Figure 1A). Plasma allantoin levels behaved almost in parallel with UA, showing significant sex differences between MFS mice but not between WT and MFS females, most likely due to the great variability of the measurements (Figure 1B). Considering sex differences in UA and allantoin plasma levels, we decided to continue working only with males because they are the most referentially studied sex for aortopathy progression as MFS patients have a significantly larger aortic root diameter than female patients. This phenotypic difference in aortic diameter between males and females is also recapitulated in MFS mice models. Moreover, male mice show minimal or no oestrogen modulation, which could additionally impact on results in females, and there were no basal differences either in UA or allantoin plasma levels between WT and MFS animals at the starting point of the study.

**Figure 1.**
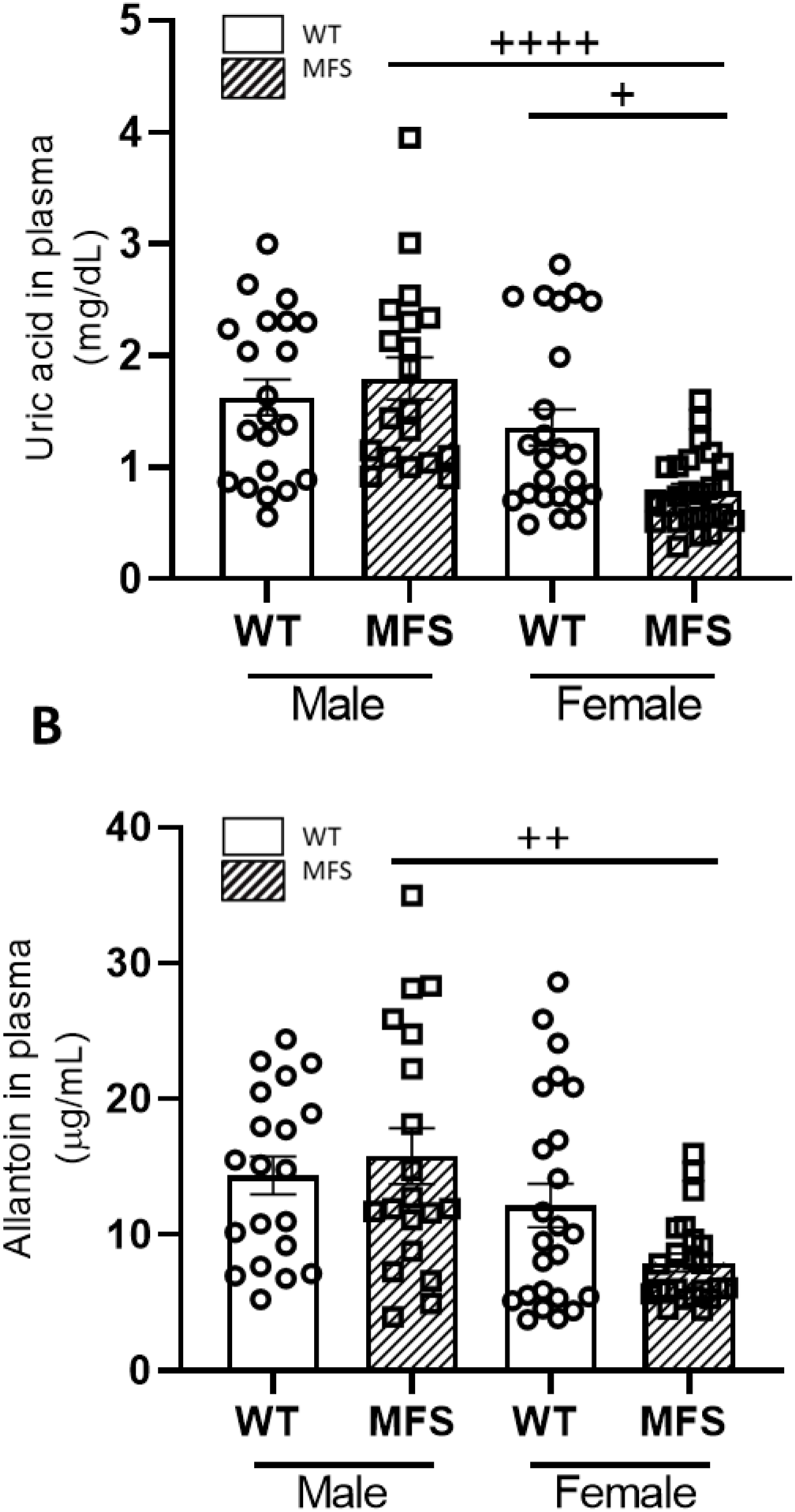
Basal uric acid and allantoin blood plasma levels in male and female WT and Marfan syndrome mice. (A) Uric acid and (B) allantoin plasma levels in two-month-old WT and MFS mice (males and females). Results are the mean ± SEM. Statistical tests: Kruskal-Wallis, Dunn’s *post-hoc*. ^+^*p* ≤0.05, ^++^*p* ≤0.01; ^++++^*p* ≤0.0001.

#### 2.2. Potassium oxonate transiently increased blood plasma levels of uric acid both in WT and MFS mice

Unlike humans and higher primates, rodents quickly catabolise UA to allantoin, which makes it difficult to study the impact of UA on the onset and/or progression of cardiovascular diseases using murine models. This problem can be transiently overcome with the administration of uricase inhibitors. Oxonic acid is the most efficient chemical inhibitor of uricase, administered as potassium oxonate (PO). Therefore, to study the impact of UA on MFS aortopathy and cardiopathy, we next generated an MFS murine model of hyperuricaemia with regular injections of PO. PO was administered to two-month-old WT and MFS mice for a period of 4, 8, and 16 weeks until three, four, and six months of age, respectively (supplementary Figure 1). After four weeks of PO treatment, plasma UA levels increased significantly in both male WT and MFS mice (treatment; *p*=0.0016), their respective increases being highly similar and, therefore, not significant between them (interaction genotype/treatment *p*=0.9737) (Figure 2A). Strikingly, plasma allantoin levels exhibited highly similar behaviour to UA, being statistically significant for PO treatment (*p* ≤0.0001) but not for the interaction genotype/treatment (*p*=0.2966) (Figure 2B). Some treated and untreated WT and MFS animals were euthanised for histopathological analysis of aortae (see below) and the rest continued PO treatment for 4 and 12 weeks more, receiving a total of 8 and 16 weeks of PO treatment, the mice reaching the age of four and six months, respectively. Note that eight-week PO treatment maintained high plasma levels of both UA and allantoin compared with untreated male mice, which only reached statistical significance in WT mice most likely due to the large variability of values obtained in the other experimental groups (Supplementary Figures 3A and B). Following a further eight weeks of PO administration in these mice (16 weeks of treatment), UA and allantoin plasma levels showed no differences between treated and untreated animals, regardless of the genotype (Supplementary Figure 4). Therefore, the PO-induced hyperuricaemia in WT and MFS mice is transient but with an effective temporal window of eight weeks after PO injection (from two to four months of age), which, nonetheless, is long enough to evaluate the potential cardiovascular impact on both WT and MFS mice.

**Figure 2.**
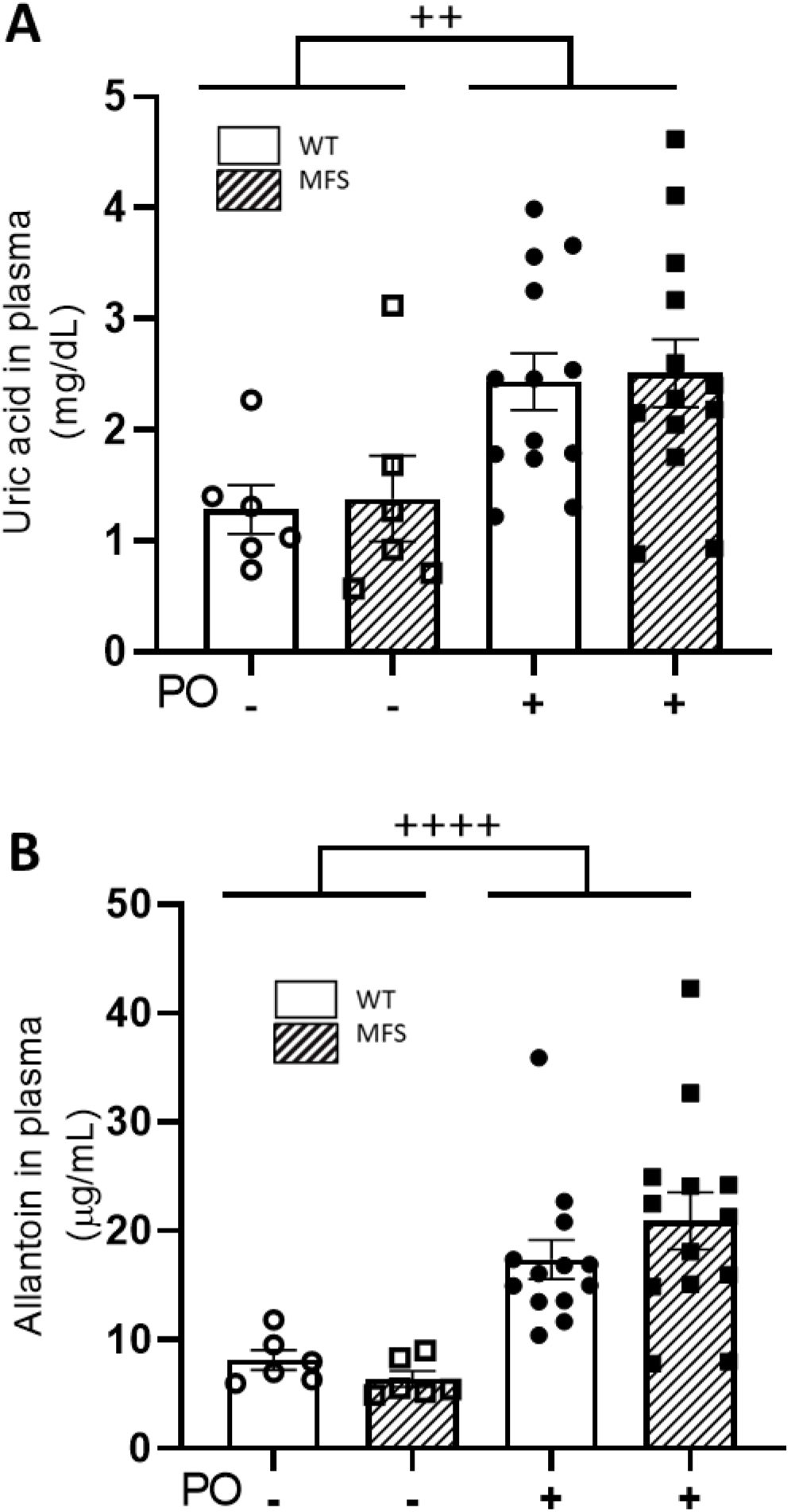
Uric acid and allantoin plasma levels following four weeks of potassium oxonate treatment. Two-month-old male WT and MFS mice were treated for four weeks with potassium oxonate (PO); subsequently, uric acid (A) and allantoin (B) plasma levels were determined. PO treatment significantly increased their respective plasma levels, however, no differences were obtained between genotypes (WT *vs.* MFS). Results are the mean ± SEM. Statistical test: Two-way ANOVA. ^++^*p* ≤0.01; ^++++^*p* ≤0.0001

#### 2.3. Aortopathy in MFS mice progressed regardless of the potassium oxonate-induced hyperuricaemia

Just before blood extraction to measure UA and allantoin levels in each PO-treated and untreated WT and MFS mouse, we analysed the aortic root diameter and cardiac parameters by 2D echocardiography. In three-month-old mice (four weeks of PO treatment), the aortic root diameter in MFS mice exhibited the characteristic aneurysm (WT/PO-*vs.* MFS/PO-) but showed no differences with PO treatment (WT/PO+ *vs.* MFS/PO+) (Figure 3). Similar results were observed when some WT and MFS animals continued receiving PO for an additional 4 or 12 weeks, reaching statistical differences between them. As expected, untreated MFS mice tended towards aortic dilatation compared with untreated WT mice (as expected) but not reaching significance due to the low number of measurements (Supplementary Figures 5 and 6).

**Figure 3.**
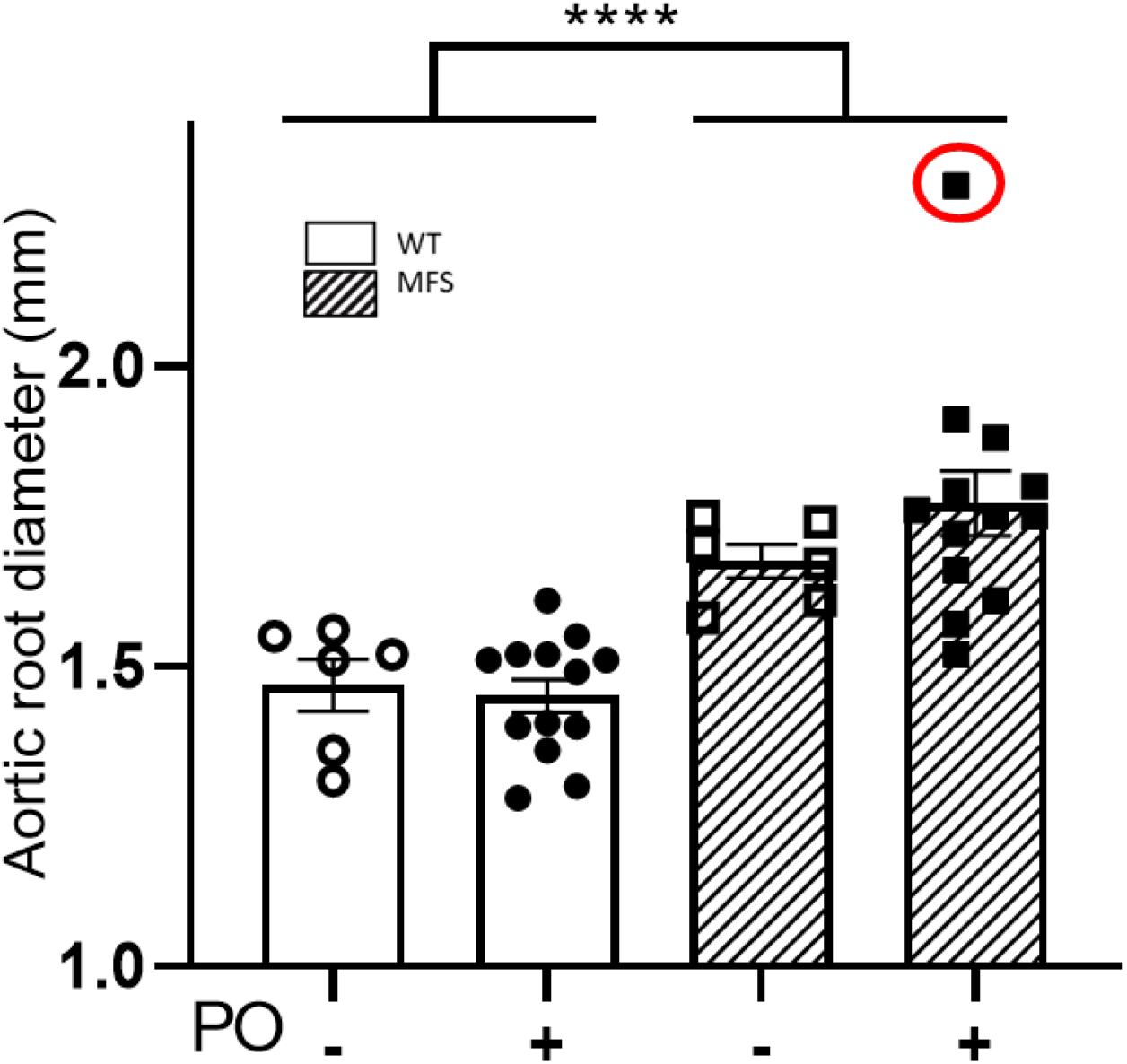
Aortic root diameter in WT and MFS mice following four weeks of potassium oxonate treatment. The aortic root diameter was measured via 2D echocardiography in two-month-old male WT and MFS mice treated for four weeks with potassium oxonate (PO). Male MFS mice show a significant increase in aortic root diameter compared with WT mice (genotype; \*\*\*\**p* ≤0.0001); however, PO-treated WT and MFS mice showed no additional differences with treatment (*p*=0.6457). An identified outlier is indicated with a red circle but was not included in the statistical analysis. Results are the mean ± SEM; Statistical test: Two-way ANOVA. \*\*\*\**p* ≤0.0001.

The echocardiographic analysis of different cardiac parameters (IVSD, LVPWD, LVDD, and LVDS) did not show any significant difference between WT and MFS mice, regardless of the length of PO treatment (Table 1).

**Table 1.** Cardiac parameters measured by 2D ultrasound in WT and MFS treated (+) or not (-) with potassium oxonate (PO; in weeks).

|  |  | WT | WT | WT | WT | MFS | MFS | MFS | MFS |
| --- | --- | --- | --- | --- | --- | --- | --- | --- | --- |
| Potassium<br>oxonate<br>(PO) | Mice age<br>(months)/<br>PO<br>(weeks) | IVSD | LVPWD | LVDD | LVDS | IVSD | LVPWD | LVDD | LVDS |
| ----- | 2 m/0 w | 0,82±0,01 | 0,79±0,01 | 1,98±0,02 | 1,27±0,03 | 0,81±0,01 | 0,77±0,01 | 1,98±0,03 | 1,25±0,03 |
| - | 3 m/4 w | 0,80±0,02 | 0,76±0,03 | 2,07±0,11 | 1,27±0,04 | 0,80±0,02 | 0,78±0,02 | 2,06±0,04 | 1,36±0,04 |
| - | 4 m/8w | 0,87±0,04 | 0,95±0,09 | 1,88±0,05 | 1,29±0,08 | 0,95±0,09 | 0,95±0,09 | 2,31±0,19 | 1,39±0,05 |
| - | 6 m/16w | 0,87±0,03 | 0,85±0,03 | 2,05±0,07 | 1,40±0,04 | 0,87±0,01 | 0,83±0,02 | 2,01±0,03 | 1,32±0,01 |
| + | 3 m/4 w | 0,83±0,01 | 0,80±0,002 | 2,03±0,05 | 1,38±0,02 | 0,82±0,01 | 0,81±0,01 | 2,07±0,06 | 1,41±0,04 |
| + | 4 m/8 w | 0,84±0,04 | 0,93±0,02 | 2,21±0,19 | 1,43±0,07 | 1,01±0,05 | 1,06±0,02 | 2,52±0,07 | 1,71±0,08 |
| + | 6 m/16 w | 0,79±0,01 | 0,80±0,02 | 2,11±0,06 | 1,35±0,04 | 0,79±0,02 | 0,75±0,03 | 2,00±0,06 | 1,34±0,02 |

#### 2.4. The characteristic structural disarray of MFS aorta persisted following potassium oxonate-induced hyperuricaemia.

We next analysed the structural organisation of the aortic wall in four- and six-month-old mice who received PO treatment for 4 and 16 weeks, respectively (Figures 4A and B). The tunica media of MFS aorta showed numerous characteristic ruptures of elastic laminae, whose extension and number were rather variable but always greater than in WT aorta. The large number of elastic breaks observed in PO-treated MFS mice aortae was indistinguishable from untreated MFS animals, regardless of the weeks of treatment (Figure 4A/4 weeks PO and 4 months of age mice and 4B/16 weeks PO and 6 months of age mice).

**Figure 4.**
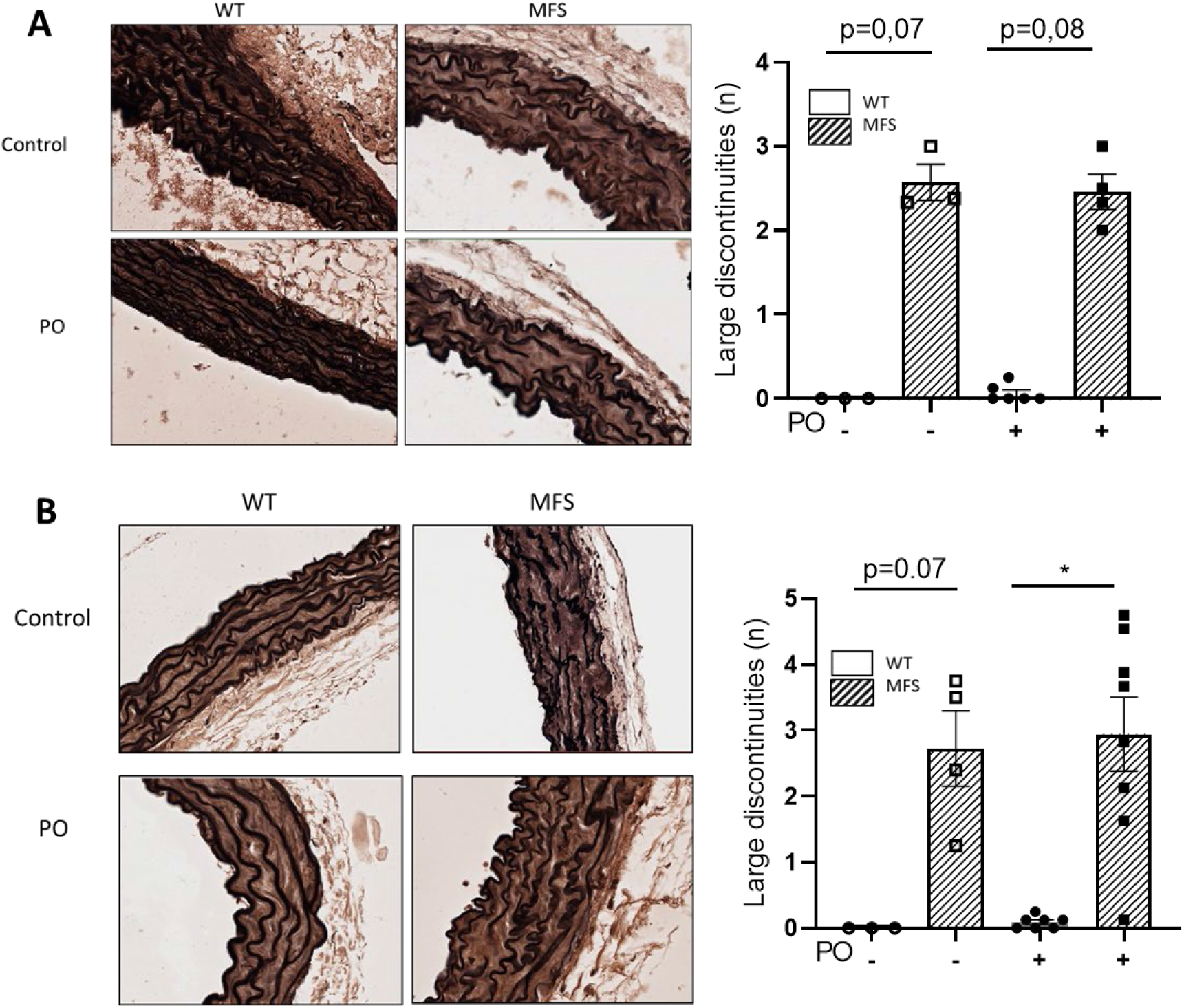
Aortic wall organisation in WT and MFS mice treated with potassium oxonate. Representative light microscope images of elastin histological staining (Elastin Verhoeff-Van Gieson) of aortic paraffin sections of the tunica media of the ascending aorta from WT and MFS mice treated or not with potassium oxonate (PO). The quantitative analysis of aortic elastic breaks after 4 and 16 weeks of PO treatment is also shown beside their respective images. Results are the mean ± SEM. Statistical analysis: Kruskal Wallis, Dunn’s *post-hoc*. \**p* ≤0.05))

#### 2.5. Potassium oxonate-induced hyperuricaemia does not modify the increased nuclear pNRF2 levels occurring in MFS mice

The nuclear factor erythroid 2-related factor 2 (NRF2) is a key transcription factor that regulates the expression of several antioxidant defence mechanisms. Oxidative stress triggers its phosphorylation (pNRF2), being subsequently translocated to the nucleus to activate the expression response of physiological antioxidant enzymes. Thus, we next evaluated the nuclear presence of pNRF2 in aortic paraffin sections from WT and MFS mice treated with PO for 4 and 16 weeks (Figures 5A and B, respectively). Aortic media showed a higher presence of nuclear pNRF2 in MFS than in WT smooth muscle cells, as previously observed [22], which is demonstrative of the MFS aorta suffering redox stress and cells consequently triggering the endogenous antioxidant response. However, after the administration of PO, this response did not change (Figure 5).

**Figure 5.**
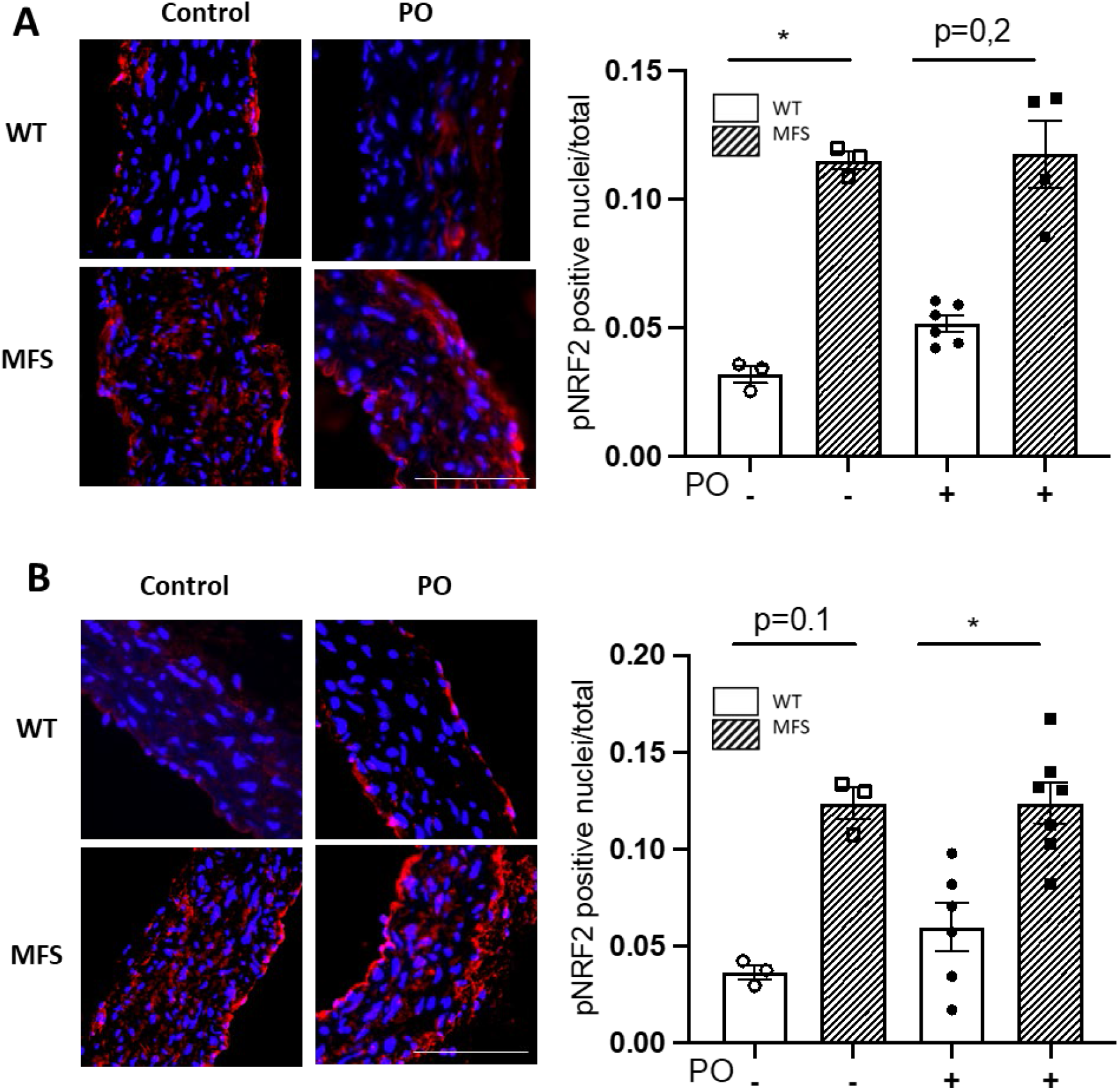
pNRF2 levels in WT and MFS mice treated with potassium oxonate. Representative images and quantitative analysis of the nuclear translocation of the phosphorylated form of NRF2 in VSMCs of the tunica media of WT and MFS mice treated with potassium oxonate (PO) for four weeks (mice aged two to three months) (A) and 16 weeks (mice aged two to six months) (B). Bar, 100 µm. Statistical tests: Kruskal-Wallis and Dunn’s multiple comparison tests. \**p* ≤ 0.05.

#### 2.5. Blood pressure is not perturbed by potassium oxonate administration

We also evaluated whether PO treatment might affect systolic blood pressure. This was measured after eight weeks of treatment (four-month-old mice) and presented no alteration (Supplementary Fig. 7).

## 3. DISCUSSION

In humans, the role of UA associated with oxidative stress and cardiovascular diseases is under debate. Most evidence comes from epidemiologic studies suggesting that increased serum levels of UA are a risk factor for cardiovascular diseases, knowing that oxidative stress is a relevant pathological mechanism in their pathogenesis and/or progression [62,63]. Allopurinol, a specific inhibitor of XOR that reduces serum levels of UA in humans, becomes protective in some cardiovascular diseases where oxidative stress is persistent (e.g., ischemia-reperfusion heart injury) [64]. Recently, we reported that allopurinol halts aortic aneurysm in a murine model of MFS, acting as an antioxidant both directly (ROS scavenger and XOR inhibitor) and indirectly by reducing the expression of metalloproteinases and NADPH oxidase Nox4. We also realised that this antioxidant action was independent of UA as plasma levels did not change throughout several treatments [22].

In humans, UA is considered the main antioxidant of the organism and, thus, it has been proposed as a potential treatment for some pathologies that affect the central nervous system [46, 65]. However, this approach in humans cannot be easily conducted and, therefore, it is necessary to first test this hypothesis in animal models. Nevertheless, in rodents, purine catabolism does not end with UA formation and, consequently, in its potential pathological accumulation (hyperuricaemia) either in serum or tissues. Hence, to study the relationship between UA and aortic aneurysm pathogenesis in a murine MFS model, we had to experimentally produce hyperuricaemia with oxonic acid, an inhibitor of uricase activity. Despite this intrinsic limitation, some hyperuricaemic mouse models have been generated that inhibit uricase, complemented or not with purine metabolism precursors where XOR is determinant for UA formation [66-69]. This pharmacological approach has been successfully used to study the impact of UA on cardio- and cerebrovascular diseases but not aortopathies [70,71]. We generated a hyperuricaemic male MFS mouse model following PO administration, and the main results obtained were the following: (i) despite aortic aneurysm occurring in MFS mice, their plasma UA levels are not significantly different from WT animals; (ii) oxonic acid, supplied as PO, produces a transient but significant elevation in plasma UA levels lasting long enough to evaluate its impact on aortopathy and cardiopathy progression in the MFS mouse model; (iii) PO-induced UA accumulation in blood plasma did not impact on either aortic aneurysm progression or the cardiac parameters examined in the MFS mouse model; and (iv) PO-induced hyperuricaemia does not modify the characteristic cellular anti-redox stress response occurring in MFS aortic media revealed by the nuclear translocation of pNRF2 in aortic vascular smooth muscle cells (VSMCs).

One striking observation was that plasma allantoin levels did not decrease in parallel with the increase in UA given that PO inhibits uricase activity. We think that this can be explained by the known non-enzymatic transformation of UA into allantoin. Thus, UA can react with anion superoxide, become oxidised and converted into allantoin. Likewise, UA can also be transformed into allantoin under alkaline conditions [72]. The overaccumulation of UA might explain the high allantoin levels.

Despite evidence supporting a role for RS in human aortic aneurysms, a clear relationship between systemic RS, serum UA levels, total antioxidant capacity, and ascending aortic aneurysms has not been definitively established. Nonetheless, serum UA concentrations and total antioxidant capacity have been reported to be associated with aortic dilatation in humans, suggesting serum UA levels as an indicator of oxidative stress [23,50, 51,73-75]. This conclusion has not been confirmed in a hyperuricaemic-like MFS mouse. However, for strictly comparative reasons and to clarify this question in humans once and for all, it would be necessary to determine serum UA levels in MFS patients over time, analysing any potential association with the progression of the aortic dilatation (absolute aortic diameter and/or Z-score). Of interest, a recent study found a highly similar protective effect of allopurinol in the same MFS mouse model used in our previous study [22], however, strikingly, they observed higher plasma UA levels in MFS mice [23]. This result, which contradicts ours reported here, is not easy to explain given that in both our studies [22 and this study], we obtained highly similar plasma UA levels measured using two different techniques (chromatographic techniques in [22] and Spinreact Uric Acid kit, this study). Moreover, we noticed that plasma UA values differed significantly depending on the blood extraction method used (infraorbital or cardiac punctures), being much higher when extracted from the heart (our unpublished observations), which is the source of measured plasma UA in [23].

We are aware that our study has some limitations. Firstly, the use of rodents to study the impact of UA, whose results are not easily translatable to humans because rodents very rarely accumulate UA. Secondly, we cannot exclude that, despite the absence of any effect on aortopathy progression in hyperuricaemic MFS mice, there could be other alterations associated with hyperuricaemia such as kidney damage [76,77] or phenotypic switching of VSMCs [78,79]. Thirdly, the use of PO generates a transient hyperuricaemic mouse model which limits the study of any potential long-term effects of hyperuricaemia. In this case, the generation of uricase (*Uox*)-deficient MFS mice would be a more suitable model. Fourthly, we only studied male mice as they have a greater aortopathy penetrance than females. Therefore, we cannot discard that females could show a different behaviour.

Based on our findings, we can conclude that, unlike ROS, UA accumulation seems not to be instrumental in the aortopathy and cardiopathy of a murine model of MFS, at least once the aneurysm is already established.

## 4. MATERIAL AND METHODS

### 4.1. Chemicals and reagents

Sodium carboxy methyl cellulose (0.5%; CMC-Na) (C5678 Sigma-Aldrich) was prepared with sterile physiological saline as the solvent for potassium oxonate (PO; 156124 Sigma-Aldrich). Potassium oxonate was suspended in CMC-Na. The injected dose was 250 mg/kg and adjusted to the weight of each animal.

### 4.2. Mice, experimental design, and study approval

Male MFS mice with a fibrillin-1 mutation *(Fbn1^C1041G/+^)* (hereafter, MFS mice) were purchased from The Jackson Laboratory (B6.129-Fbn1tm1Hcd/J; Strain #012885/Common name: C1039G; Bar Harbor, ME 04609, USA). MFS and sex- and age-matched wild-type littermates (WT mice) were maintained in a C57BL/6J genetic background. All mice were housed according to the University of Barcelona’s institutional guidelines (constant room temperature at 22°C, controlled environment 12/12-hour light/dark cycle, 60% humidity and *ad libitum* access to food and water). WT and MFS mice were randomly divided into four experimental groups: WT and MFS mice treated with CMC-Na (vehicle), and WT and MFS mice treated with PO. All four groups received the treatment by intraperitoneal injection of the corresponding reagent (CMC-Na: 0.5%; PO: 250 mg/kg) every 48 h (Supplementary Figure 1). Animal care and colony maintenance were carried out according to European Union (Directive 2010/63/EU) and Spanish guidelines (RD 53/2013) for the use of experimental animals. Ethical approval was obtained from the local animal ethics committee (CEEA protocol approval number: 357/22).

### 4.3. Uric acid and allantoin analysis in blood plasma

Plasma uric acid levels were measured using Spinreact (model Spinlab 100) with the Spinreact Uric Acid kit (SP41001). We followed the manufacturer’s instructions. For the determination of allantoin in blood plasma, an adapted protocol was used, as previously described [80]. Briefly, plasma (60 µl) was deproteinised with acetonitrile (25 µl). Samples were centrifuged (5 min, 12,000 *g*). Ten µl of supernatant was injected into the HPLC system. Separation of allantoin was performed on a Synergy Hydro-RP C-18 reversed-phase column (250 × 4.6 mm I.D., 5 m particle size) from Phenomenex (Torrance, CA, USA). Allantoin elution (at 4 min) was performed with potassium dihydrogen phosphate (10 mM, pH 2.7): acetonitrile (85:15) and ultraviolet detection (at 235 nm).

### 4.4. Echocardiography

Two-dimensional transthoracic echocardiography was performed in all animals under 1.5% inhaled isoflurane. Each animal was scanned 12–24 hours before sacrifice. Images were obtained with a 10–13 MHz phased array linear transducer (IL12i GE Healthcare, Madrid, Spain) in a Vivid Q system (GE Healthcare, Madrid, Spain). Images were recorded and later analysed offline using commercially available software (EchoPac v.08.1.6, GE Healthcare, Madrid, Spain). Proximal aortic segments were assessed in a parasternal long-axis view. The aortic root diameter was measured from inner edge to inner edge in end-diastole at the level of the sinus of Valsalva. Left ventricle (LV) dimensions were assessed in 2D mode in a parasternal long-axis view at both end-diastole (LVDD) and end-systole (LVSD). The interventricular septum and posterior wall thickness at end-cardiac diastole were also measured. All echocardiographic measurements were carried out in a blinded manner by two independent investigators with no knowledge of genotype or treatment.

### 4.5. Histopathology

Paraffin-embedded tissue arrays of mice aortae from different experimental sets were cut into 5 µm sections. Elastic fibre ruptures were quantified by counting the number of large fibres breaks in tissue sections stained with Verhoeff-Van Gieson. Breaks larger than 20 µm were defined as evident large discontinuities in the normal circumferential continuity (360°) of each elastic lamina in the aortic media [81]. They were counted along the length of each elastic lamina in four different representative images of three non-consecutive sections of the same ascending aorta. Three sections per condition were usually studied, spaced 10 µm apart (two sections). All measurements were carried out in a blinded manner by two different observers with no knowledge of genotype and treatment. Images were captured using a Leica DMRB microscope (40x oil immersion objective) equipped with a Leica DC500 camera and analysed with Fiji Image J Analysis software.

### 4.6. Immunohistochemistry and immunofluorescence staining

For pNRF2 immunofluorescence, paraffin-embedded aortic tissue sections (5 μm thick) were deparaffinised and rehydrated prior to unmasking the epitope. Sections were treated first with heat-mediated retrieval solution (1 M Tris-EDTA, 0.05% Tween, pH 9) for 30 min in the steamer at 95°C. Next, sections were incubated for 20 minutes with ammonium chloride (NH^4^Cl, 50 mM, pH 7.4) to block free aldehyde groups, followed by a permeabilisation step using 0.3% Triton X-100 for 10 min and then treated with 1% BSA blocking buffer solution for 2 h prior to overnight incubation with monoclonal anti-pNRF2 (1:200; Abcam ab76026) in a humidified chamber at 4°C. On the next day, sections were rinsed with PBS followed by 60 min incubation with the secondary antibody goat anti-rabbit Alexa 647 (1:1.000, A-21246, Invitrogen). Sections were counterstained with DAPI (1:10.000) and images were acquired using an AF6000 widefield fluorescent microscope. For quantitative analysis, four areas of each ascending aorta section were quantified with Image J software. All measurements were carried out in a blinded manner by two independent investigators.

### 4.7. Blood pressure measurements

Systolic blood pressure measurements were acquired by the tail-cuff method and using the Niprem 645 non-invasive blood pressure system (Cibertec, Madrid, Spain). Mice were positioned on a heating pad and all measurements were carried out in the dark to minimise stress. All animals were habituated to the tail cuff by daily training one week prior to the final measurements. Then, the systolic blood pressure was recorded over the course of three days. For quantitative analysis, the mean value of three measurements per day was used per animal. All measurements were carried out in a blinded manner with no knowledge of genotype or experimental group.

### 4.8. Statistical analysis

Data were presented as bars showing the mean ± standard error of the mean (SEM). Firstly, normal distribution and equality of error variance data were verified with Kolmogorov-Smirnov/Shapiro-Wilk tests and Levene’s test, respectively, using the IBM SPSS Statistics Base 22.0 before parametric tests were used. Differences between three or four groups were evaluated using one-way or two-way ANOVA with Tukey’s *post-hoc* test if data were normally distributed and variances were equal, or the Kruskal-Wallis test with Dunn’s *post-hoc* test if data were not normally distributed. For the comparison of two groups, the unpaired t-test was utilised when data were normally distributed and variances were equal, or the Mann-Whitney U test if data did not follow a normal distribution. A value of *P* ≤0.05 was considered statistically significant. Data analysis was carried out using GraphPad Prism software (version 9.1.2; GraphPad Software, La Jolla, CA). Outliers (ROUT 2%, GraphPad Prism software) were removed before analysis.

## Supplementary Materials

Figure S1: Schematic representation of the experimental protocols followed in this study.

Figure S2: Aortic root diameter in WT and MFS mice at the start time of the study (two months old).

Figure S3: Uric acid and allantoin plasma levels following eight weeks of treatment with potassium oxonate.

Figure S4: Uric acid and allantoin plasma levels following 16 weeks of treatment with potassium oxonate.

Figure S5: Aortic root diameter in WT and MFS mice treated with PO for eight weeks.

Figure S6: Aortic root diameter in WT and MFS mice treated with PO for 16 weeks.

Figure S7: Blood pressure in WT and MFS treated with potassium oxonate.

## Author Contributions

Conceptualisation, G.E.; methodology, I.R-R. A.L-S., M.P-B., B.P.; formal analysis, I.R-R, A.L.-S., V.C., G.E.; investigation, I.R.-R. A.L.-S., M.P-B. B.P., F.J-A., V.C., G.E.; resources, I.R.-R.; data curation, F.R-R, V.C., G.E.; writing— original draft preparation, G.E.; writing—review and editing, I.R-R., F.J-A, V.C., G.E.; supervision, F.J-A., V.C., G.E.; funding acquisition, F.J.-A., V.C., G.E. All authors have read and agreed to the published version of the manuscript.

## Institutional Review Board Statement

The animal study protocol was approved by the local Committee of Ethical Animal Experimentation (CEEA-PRBB; Protocol Number: 357/22) in accordance with the guidelines of the European Communities Directive 86/609/EEC. The PRBB has Animal Welfare Assurance (#A5388-01, Institutional Animal Care and Use Committee approval date 05/08/2009), granted by the Office of Laboratory Animal Welfare (OLAW) of the US National Institutes of Health.

Informed Consent Statement: Not applicable.

## Data Availability Statement

Original data can be requested from the corresponding author.

## Acknowledgements

We thank Fernando J. Pérez Asensio (Animal Room of the PCB, Barcelona) for uric acid measurements and Helena Kruyer for English editorial assistance.

## SUPPLEMENTARY FIGURE LEGENDS

**Supplementary Figure 1.**
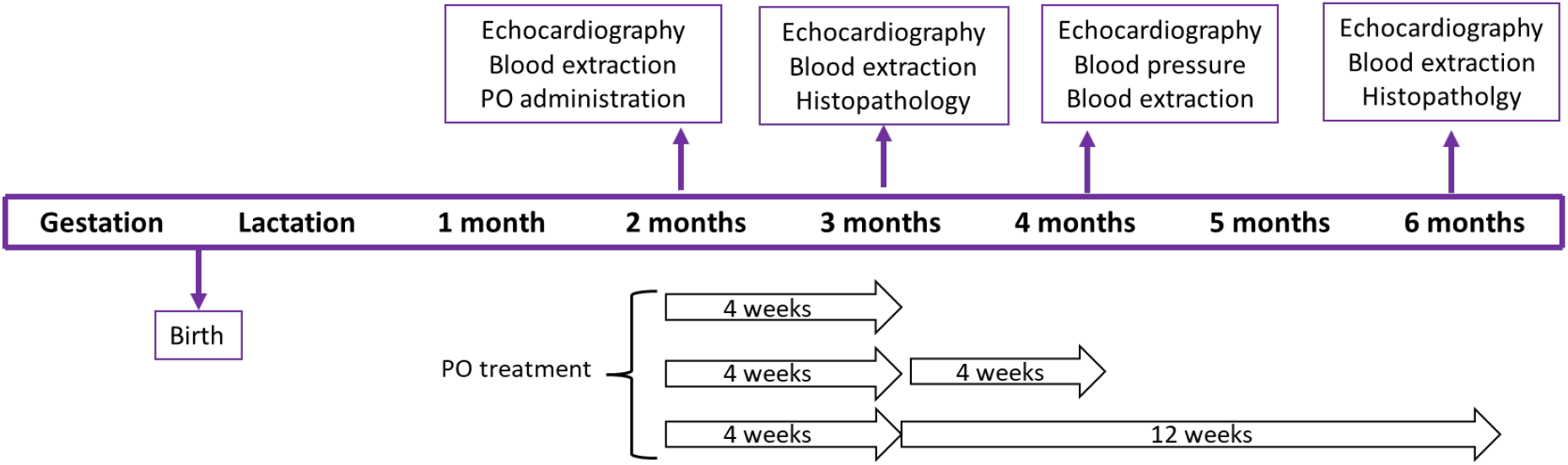
Representative scheme of the experimental protocols for allopurinol treatments. PO: potassium oxonate. PO was administered to WT and MFS mice for four weeks, some animals were euthanized and other continued receiving PO for additional 4 weeks or 12 weeks for a total time of 8 and 16 weeks, respectively. Inside boxes it is indicated the applied techniques.

**Supplementary Figure 2.**
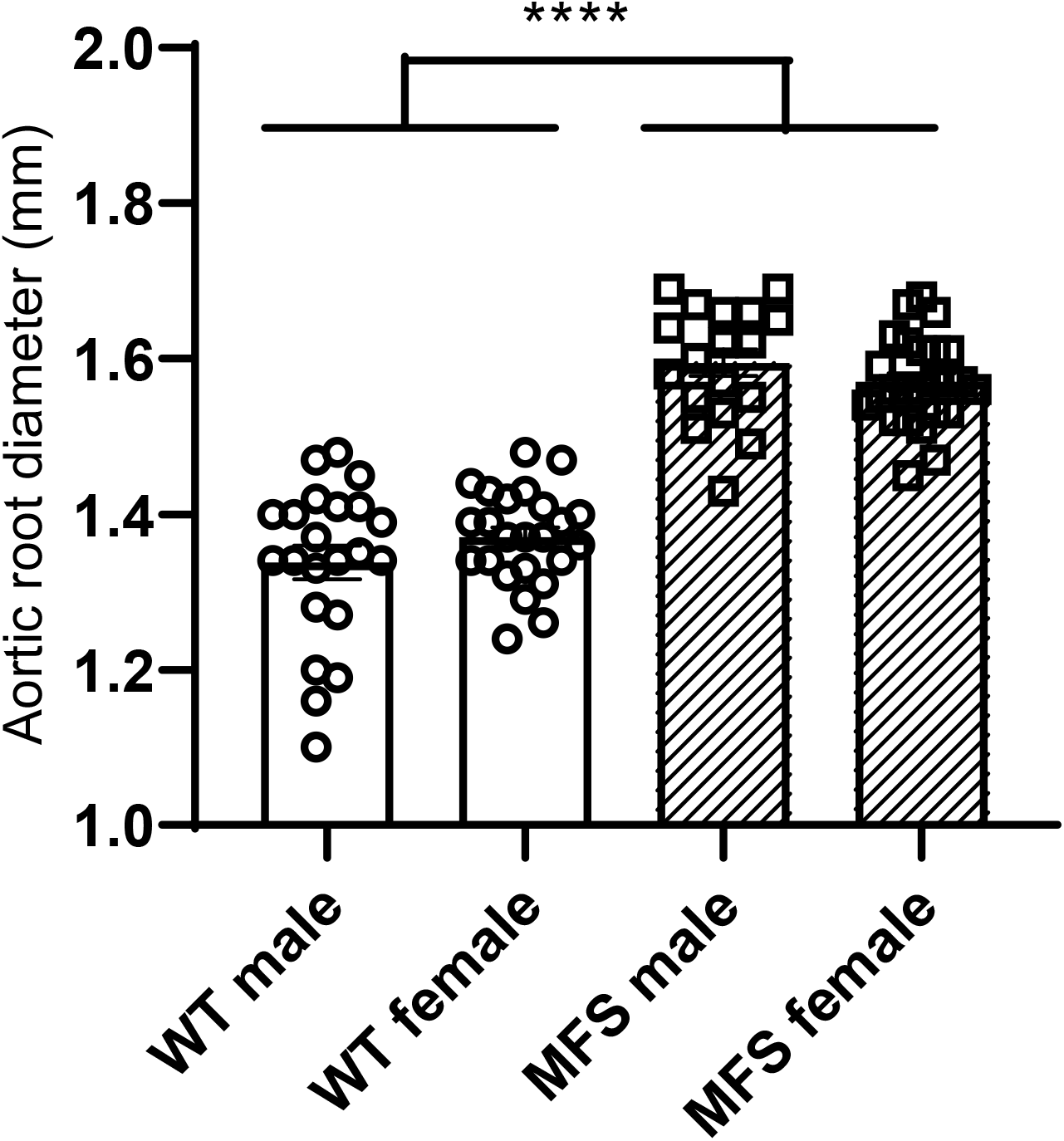
Aortic root diameter in WT and MFS mice at the start point of the study (two months old). The aortic root diameter of male and female WT and MFS mice was measured with 2D echocardiography. Both male and female MFS mice presented the expected increased aortic root diameter compared with WT mice. Results are the mean ± SEM. Statistical test: Two-way ANOVA. \*\*\*\**p* ≤0.0001.

**Supplementary Figure 3.**
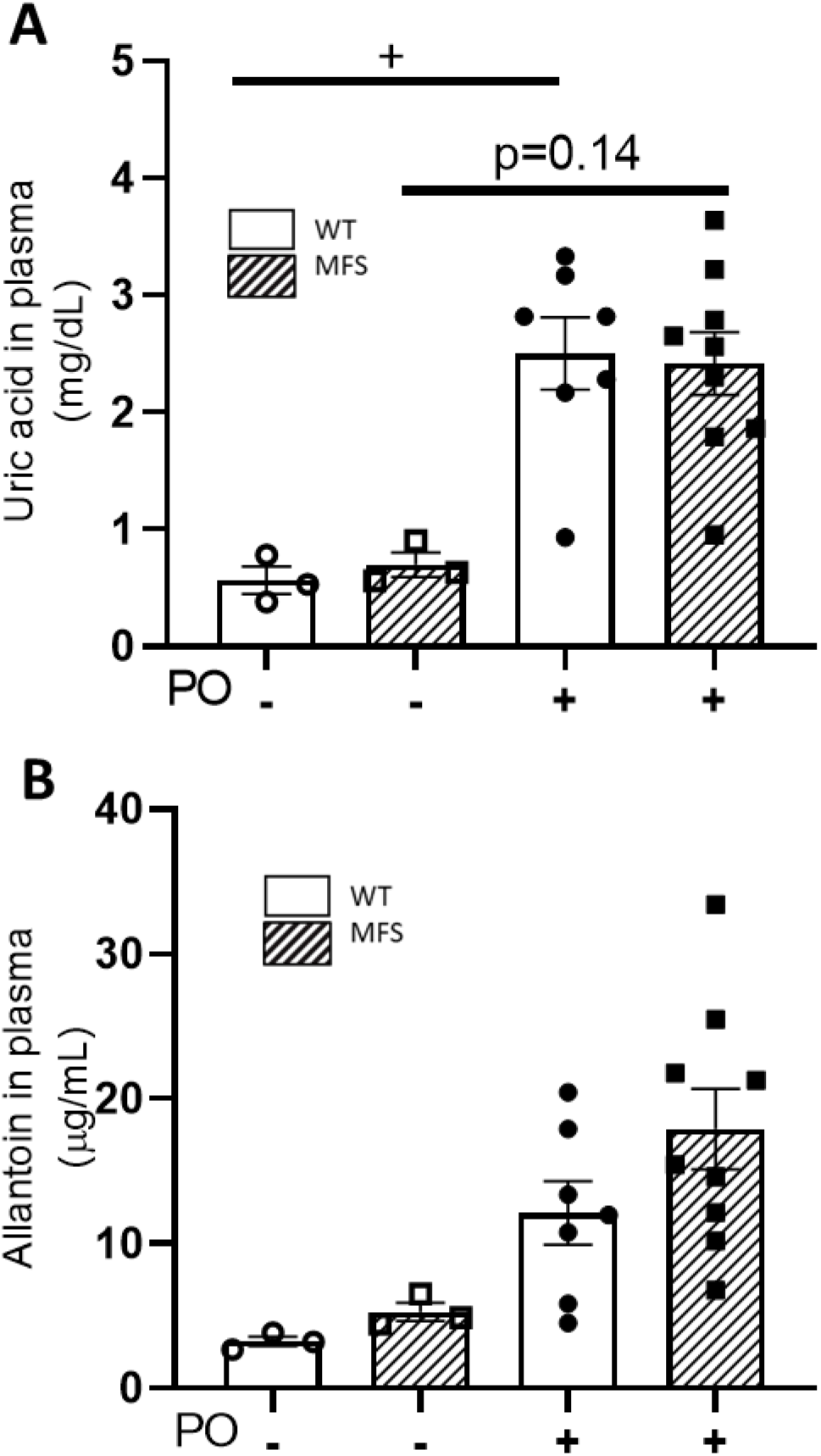
Uric acid and allantoin plasma levels following eight weeks of potassium oxonate treatment. Three-month-old WT and MFS mice (treated with PO for four weeks) continued receiving PO treatment for four weeks more (total of eight weeks of PO treatment; four-month-old mice) and their uric acid (A) and allantoin (B) plasma levels were determined. UA and allantoin plasma levels remained increased both in WT and MFS mice, however, only WT mice reached statistical significance. Results are the mean ± SEM. Statistical tests: Kruskal Wallis, Dunn’s *post-hoc*. +*p* ≤0.05

**Supplementary Figure 4.**
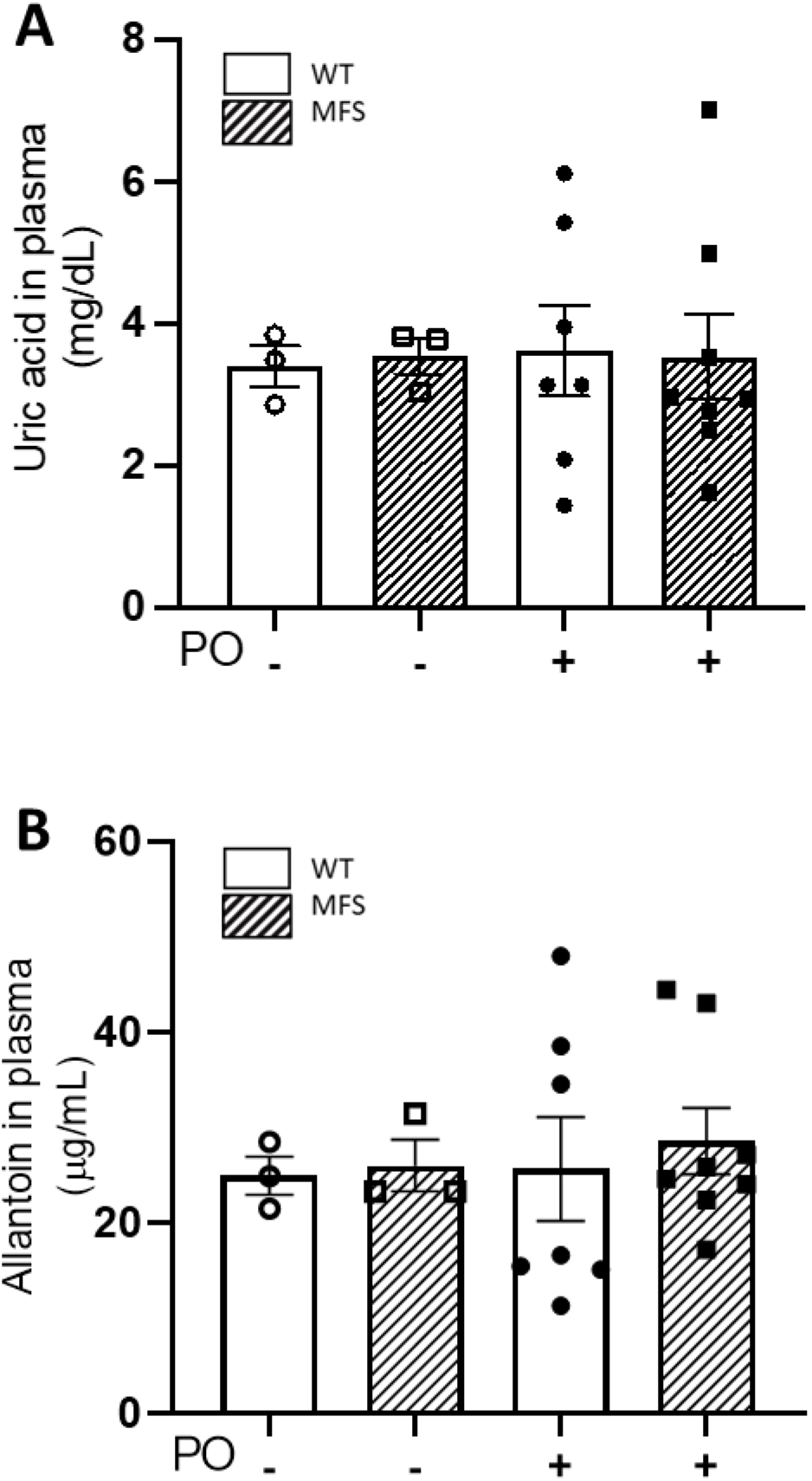
Uric acid and allantoin plasma levels following 16 weeks of potassium oxonate treatment. Male WT and MFS mice treated for eight weeks with potassium oxonate (PO) were treated for an additional eight weeks (16 weeks of PO treatment; six-month-old mice) and uric acid (A) and allantoin (B) plasma levels were determined. This longer PO treatment did not maintain the increased UA and allantoin levels obtained with the shorter treatments. Results are the mean ± SEM. Statistical test: Kruskal-Wallis.

**Supplementary Figure 5.**
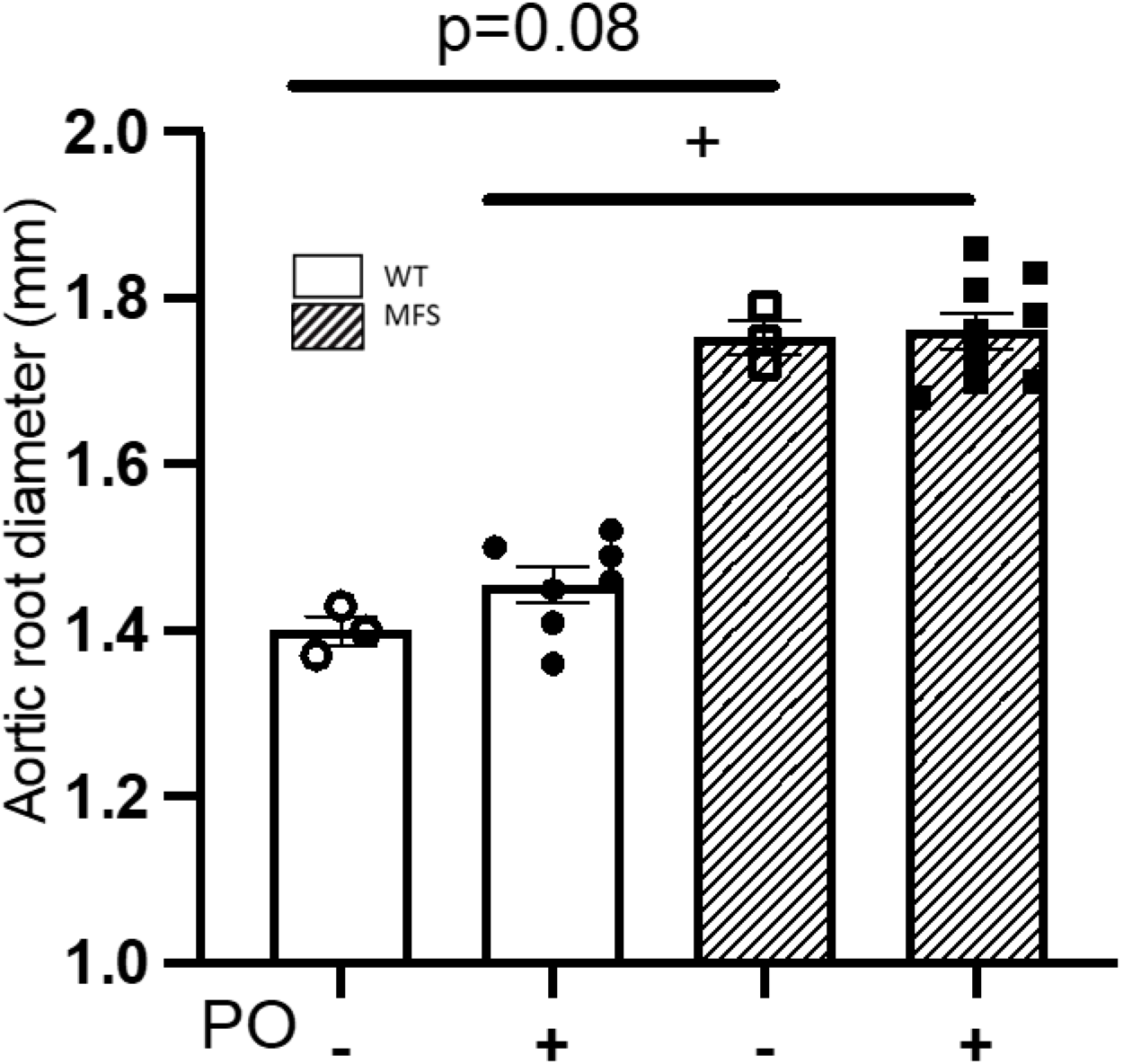
Aortic root diameter in WT and MFS mice following eight weeks of potassium oxonate treatment. Male WT and MFS mice treated for four weeks with potassium oxonate (PO) were treated for an additional four weeks (eight weeks of PO treatment; four-month-old mice) and the aortic root diameter was measured again with 2D echocardiography. MFS mice maintained the increased aortic root diameter regardless of the PO treatment. Results are the mean ± SEM. Statistical tests: Kruskal Wallis, Dunn’s *post-hoc*. \**p* ≤0.05.

**Supplementary Figure 6.**
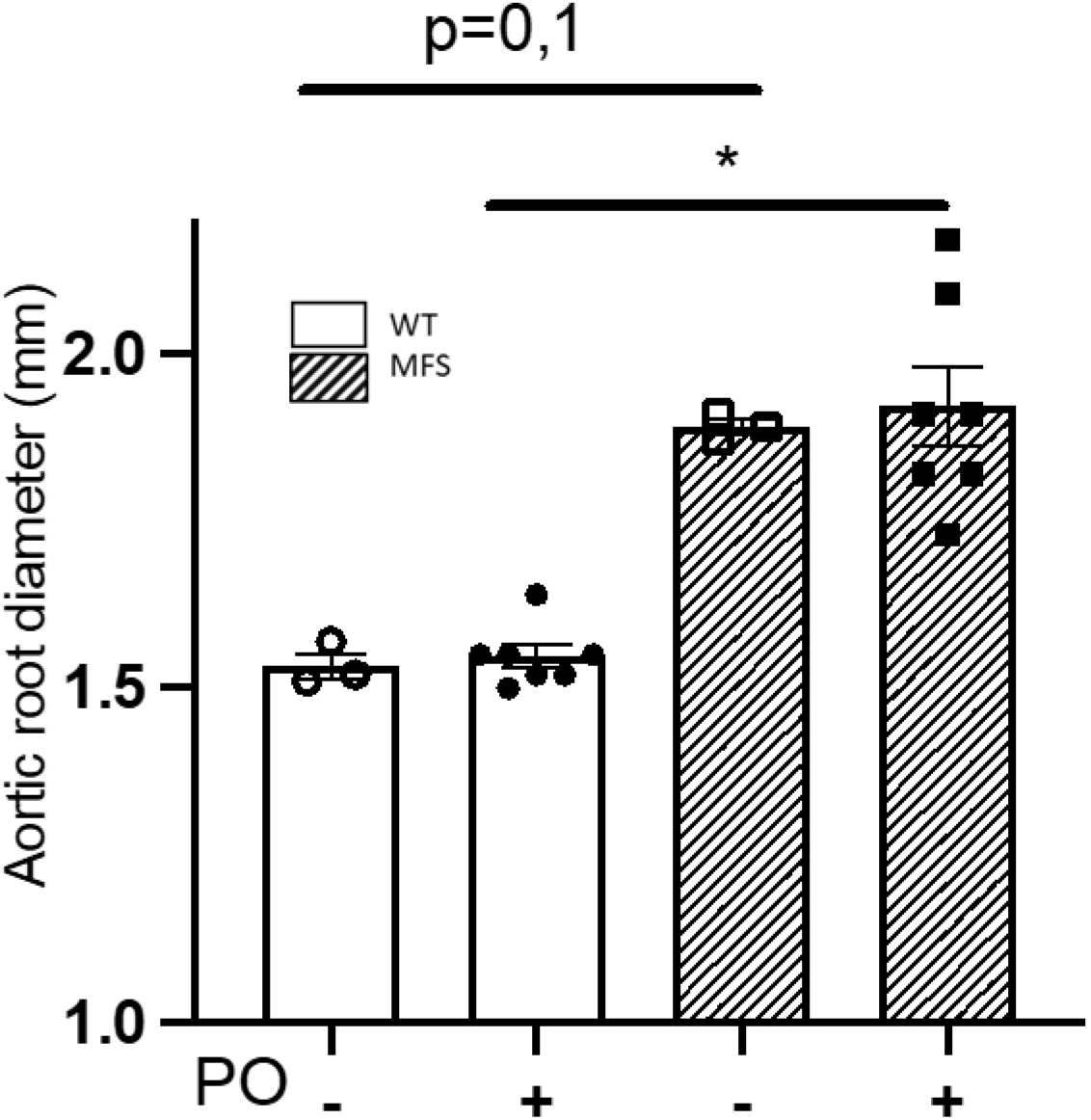
Aortic root diameter in WT and MFS mice following 16 weeks of potassium oxonate treatment. Male WT and MFS mice treated for eight weeks with potassium oxonate (PO) were treated for an additional eight weeks (16 weeks of PO treatment; six-month-old mice) and the aortic root diameter was measured again with 2D echocardiography. MFS mice maintained the increased aortic root diameter regardless of the PO treatment. Results are the mean ± SEM. Statistical tests: Kruskal Wallis, Dunn’s *post-hoc*. \**p* ≤0.05.

**Supplementary Figure 7.**
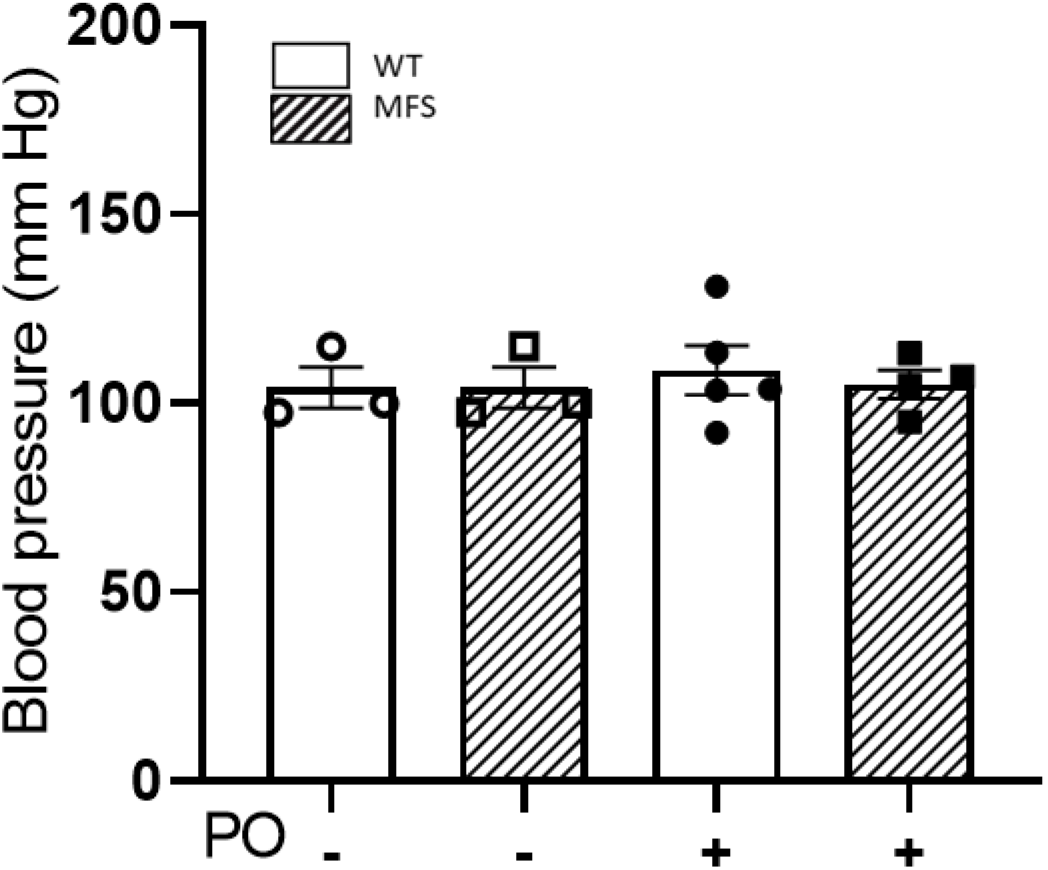
Blood pressure in WT and MFS treated with potassium oxonate. Systolic blood pressure measurements in four-month-old WT and MFS mice treated with potassium oxonate (PO; eight weeks). PO did not alter blood pressure in any experimental group of mice. Results are the mean ± SEM. Statistical test: Kruskal-Wallis with Dunn’s multiple comparison test.

